# The host-directed therapeutic imatinib mesylate accelerates immune responses to *Mycobacterium marinum* infection and limits pathology associated with granulomas

**DOI:** 10.1101/2022.11.28.518230

**Authors:** Tesia L. Cleverley, Siri Peddineni, Jeannette Guarner, Francesca Cingolani, Heather Koehler, Edward Mocarski, Daniel Kalman

## Abstract

Mycobacterial infections, including those caused by members of the mycobacterium tuberculosis complex [MTC] and Nontuberculous mycobacteria [NTM], can induce widespread morbidity and mortality in people. Mycobacterial infections cause both a delayed immune response, which limits rate of bacterial clearance, and formation of granulomas, which contain bacterial spread, but also contribute to lung damage, fibrosis, and morbidity. Granulomas also limit access of antibiotics to bacteria, which may facilitate development of resistance. MTC members resistant to some or all antibiotics are estimated to account for a third of deaths from tuberculosis [TB], and newly developed antibiotics have already engendered resistance, pointing to the need for new therapeutic approaches. Imatinib mesylate, a cancer drug used to treat chronic myelogenous leukemia [CML] that targets Abl and related tyrosine kinases, is a possible host-directed therapeutic [HDT] for mycobacterial infections, including TB. Here, we use the murine *Mycobacterium marinum* [Mm] infection model, which forms quantifiable granulomas on the tails, in conjunction with transcriptomic analysis of the tail lesions. The data indicate that imatinib induces gene signatures indicative of immune activation at early time points post infection that resemble those seen at later ones, suggesting that imatinib accelerates but does not substantially alter anti-mycobacterial immune responses. Moreover, focusing on the TNFα pathway, which is induced by imatinib, we show that imatinib promotes cell survival in infected bone marrow-derived macrophages [BMDMs] in a manner that depends on caspase 8. Moreover, imatinib limits formation and growth of granulomas, an effect abrogated in mice lacking caspase 8. These data provide evidence for the utility of imatinib as an HDT for mycobacterial infections in accelerating immune responses, and limiting pathology associated with granulomas, and thus mitigating post-treatment morbidity.

**Author Summary:** Mycobacterial infections remain an important cause of morbidity and mortality in humans; for example, *Mycobacterium tuberculosis* [Mtb], the cause of tuberculosis [TB], kills ∼1.5 million and newly infects ∼10 million each year. Although most people effectively combat mycobacterial infections, treatment is compromised in at-risk individuals by an indolent immune response and chronic inflammation, which results in granulomas that encase the bacteria and limit spread. Granulomas also contribute to tissue damage and limit access of antibiotics to bacteria, which engenders resistance. We proposed using imatinib mesylate, a host directed therapeutic [HDT], against mycobacteria. Imatinib, a cancer therapeutic that inhibits Abl and related tyrosine kinases, alters intracellular transit of bacteria during infection. Using systems biology approaches in conjunction with murine infections with *Mycobacterium marinum*, a close genetic relative of Mtb that forms tail granulomas, we report that imatinib does not fundamentally alter the anti-mycobacteria immune response, but rather accelerates it. In addition, imatinib limits granuloma formation and growth, an effect abrogated in mice lacking caspase 8. These data highlight imatinib as a possible HDT for mycobacterial infections including TB with the capacity to augment the immune response in at-risk individuals, and limit granuloma growth, thereby limiting tissue damage.

## Introduction

Pathogenic mycobacteria have developed numerous strategies to manipulate and evade host immune responses. Notable members of this genus include *Mycobacterium tuberculosis* [Mtb], the causative agent of tuberculosis [TB], a leading cause of morbidity and mortality that claims 1.5 million lives each year (1), *Mycobacterium leprae*, which causes Hansen’s disease (leprosy), *Mycobacterium avium-intracellulare*, an opportunistic pathogen affecting immunocompromised patients and those with severe lung disease such as cystic fibrosis, and *Mycobacterium marinum* [Mm], a human pathogen acquired from contaminated aqueous environments or infected fish that causes skin lesions called “fish tank granulomas”(2). All these bacteria are either naturally resistant to antibiotics or readily acquire resistance (3, 4). The standard of care for TB, for example, is a multi-drug antibiotic regimen given over four to nine months. Importantly, Mtb strains resistant to some or all available antibiotics have emerged (5, 6), including stains resistant to newly developed antibiotics such as bedaquiline, delamanid, and pretomanid (7, 8), highlighting the need for novel treatment strategies for TB.

Mycobacteria subvert the host immune response in a variety of ways. Infection with Mtb attracts macrophages and other innate cells to the site of infection (9) but limits activation and cytokine production of antigen presenting cells (10). Within macrophages, Mm and Mtb prevent phagolysosomal fusion (11-13), which both precludes activation of macrophages and limits antigen presentation (14-16). Mtb has also been shown to limit maturation of dendritic cells (DCs) and their migration to lymph nodes (17, 18), thereby delaying the onset of adaptive responses (19-21). Accordingly, in humans adaptive immune responses to Mtb emerge approximately 42 days after exposure (22, 23). Notably mycobacterial infections are accompanied by chronic inflammation, possibly facilitated by secretion of antigenic decoy proteins such as Antigen 85 (24). Yet the immune response to mycobacteria remains highly effective in most people, and in TB, it is estimated that 90% of those infected with Mtb maintain control of the infection and do not develop active disease (1, 25). However, for all mycobacterial infections, those with immunocompromising conditions remain far more susceptible(26). An important question concerning mycobacteria treatment strategies for people with chronic disease remains how to facilitate a more efficient or efficacious immune response (24).

With a delay in immune responses and chronic inflammation, the infected host forms granulomatous lesions that appear to contain the bacteria and thereby limit its spread. Granuloma formation is in part mediated by tumor necrosis factor α [TNFα](27-31) and involves congregation of macrophages that enlarge and have more cytoplasm, known as epithelioid macrophages, which surround and ingest infected cells (32, 33). Granulomas also contain neutrophils, dendritic cells, B cells, T cells, and natural killer cells, together with fibroblasts that produce extracellular matrix [ECM](33). Epithelialioid macrophages and fibroblasts encase the infected cells and limit bacterial dissemination. Many granulomas exhibit necrosis (34, 35), which may contribute to chronic inflammation that damages surrounding lung tissue and impairs respiratory function (36). Moreover, the structure of the granuloma reduces penetrance of antibiotics, resulting in suboptimal antibiotic concentrations within the granuloma, thereby facilitating development of resistance (37, 38). Chronic inflammation results in tissue destruction that contributes to the development of fibrosis and reduces elasticity of lung tissue. As a result, half the patients successfully treated for TB exhibit lasting respiratory impairment (39-41), further exacerbating the economic burden of the disease (42). While much is known about structure and formation of granulomas, less is known about how to resolve granulomas and restore lung function in TB patients following treatment, or whether resolution would allow bacteria to escape, thereby exacerbating disease.

To address the need for novel therapeutics for mycobacterial disease, we have been developing imatinib as an adjunctive host-directed-therapeutic (HDT). Imatinib is a well-tolerated cancer drug that inhibits tyrosine kinase activity of c-Abl, c-Kit, and platelet derived growth factor receptor [PDGFR], and is the frontline therapy for chronic myelogenous leukemia [CML] and gastrointestinal stromal tumors [GISTs]. As an HDT for infections, imatinib is less likely to engender resistance compared to conventional antimicrobial drugs (43). Upon infection with Mtb or Mm infection, imatinib induces phagolysosomal fusion in mouse monocytes (44) and in human macrophages (45), an effect evident at micromolar concentrations. At low doses in mice, imatinib also induces myelopoiesis (46). Finally, prophylactic administration of imatinib also limits development of granulomas induced by Mm in mice (44). Notably, fibrosis may contribute to bacterial persistence and tissue dysfunction even after discontinuation of antibiotic chemotherapy (36, 39, 40, 47). Fibrosis in lung, skin and other tissues associated with noninfectious causes is mediated by PDGFR, also a target of imatinib, and imatinib has shown efficacy in these indications (48-53), raising the possibility that the drug may likewise limit fibrosis and granuloma-associated pathology.

The observation that imatinib limits Mm infections and granuloma formation, and induces myelopoiesis, led us to hypothesize that the drug might accelerate the formation of anti-mycobacteria immune responses, and in so doing might limit formation and/or promote resolution of granulomas. To test this idea, we chose the Mm mouse model of mycobacterial infection, in which granulomas develop on the tail within two weeks after infection. We defined gene expression signatures in tail granulomas and asked how imatinib impacts such signatures at different time points post infection. We found that imatinib accelerates appearance of gene signatures associated with immune cell activation in response to infection, including, for example, the macrophage activation marker TNFα. Finally, we found that imatinib effects on granulomas are abrogated in mice lacking caspase 8, a component of TNFα signaling pathways.

## Results

### Imatinib limits formation of granulomas in mice infected with Mm

Imatinib mesylate, administered to mice starting one day prior to IV infection with low inoculums of Mm (∼5 Log_10_ CFU/ mouse), reduced CFUs in the tail, spleen, or lung, whereas no reduction was evident in control animals provided with water as a control (44). At higher inoculums [>6 Log_10_ CFU/mouse), the drug was without effect on CFUs in any tissue at 6-, 14-, or 21-days post infection [p.i.] (Figures 1A-C). Notably, such high inoculums induce more rapid formation of granulomatous lesions on the tail, usually within 4-12 days (Figure 1D), which continue to increase in size before reaching their maximal extent by 14-21 p.i. (Figure 1E). Lesion size was quantified over time by measuring the length of each lesion on the surface of the tail to determine the change in lesion size over the treatment period (Figures 1F and G). With imatinib treatment beginning one day prior to infection with 10^7^CFU/ mouse, development of granulomatous lesions was restricted as reported (44). In an experiment using a lower inoculum (2×10^6^ CFU/mouse, Figure 1H) only 20% of imatinib-treated animals developing lesions during the first week compared to 40% of animals treated with water (Figure 1I). To determine whether imatinib limited development of actively growing lesions, the drug was administered starting at day seven p.i. and continuing until day 14 (Figure 1H). Over this time period, lesions from water-treated mice grew an average of 13mm, whereas lesions from imatinib-treated animals grew on average 6.5mm (Figure 1J, F and G). To determine effects of imatinib on established lesions, lesions were allowed to develop for 14 days prior to administration of imatinib for an additional seven days (Figure 1H). Whereas lesions on control mice grew by an average of 6.6mm during this period, no net growth was evident in mice treated with imatinib, with lesion size increasing in some animals (6 of 13 animals), but either not changing (3 of 13 animals) or decreasing (4 of 13 animals) in others (Figure 1K). Together, these data suggest that imatinib limits formation and growth of granulomas in a manner that does not depend on bacterial load.

**Figure 1:**
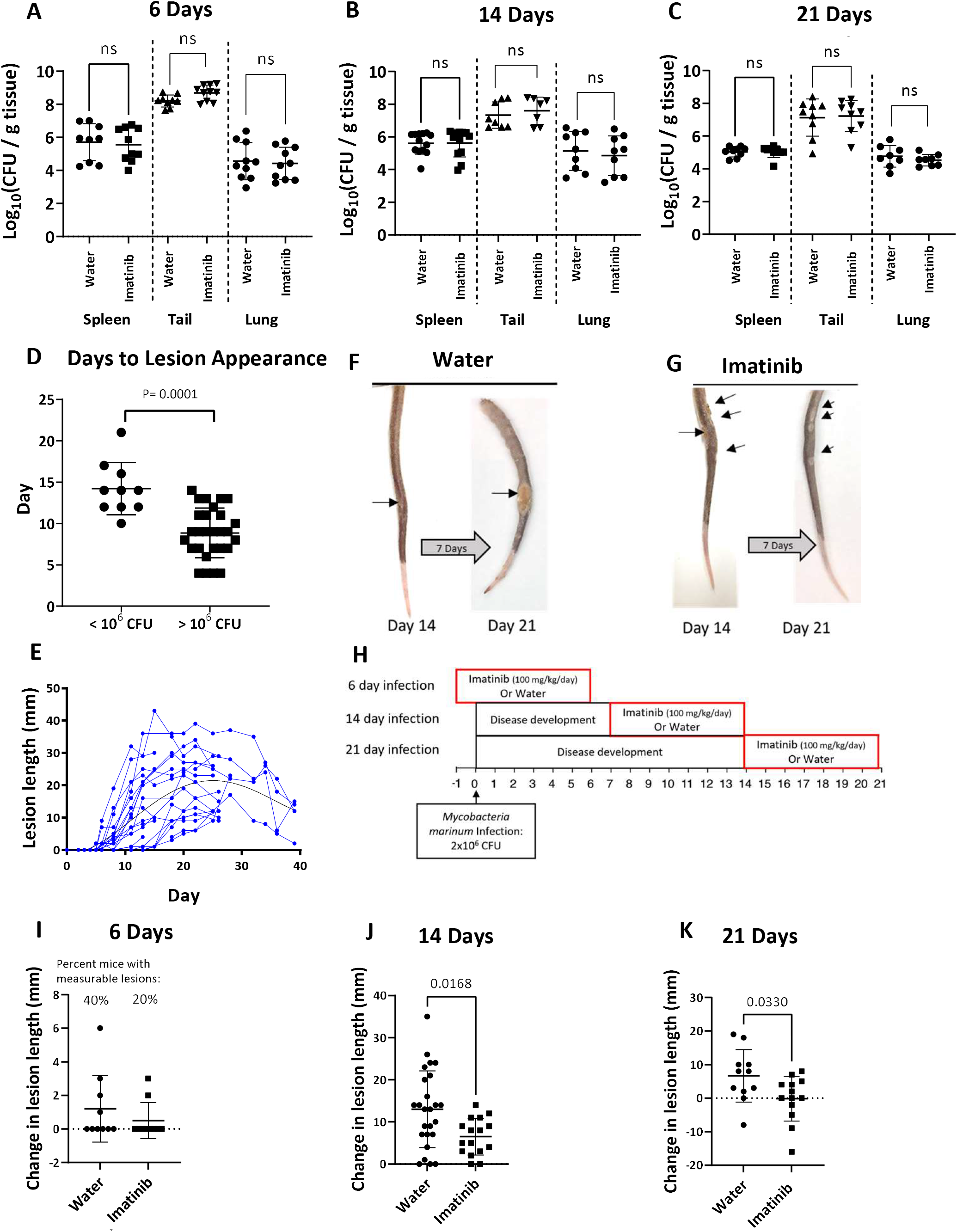
Imatinib causes a reduction in the size of granulomatous lesions in Mm-infected mice. **A-C**. Bacterial load in the spleen, tail, and lung of mice infected with 2×10^6^ CFU of Mm and treated with 100mg/kg/day imatinib via osmotic pump starting day -1, day 7, or day 14 for at total infection period of 6 days **(A)**, 14 days **(B)**, and 21 days **(C). D**. Relationship between the number of days post infection that lesions are first visible on the tails as a function of the inoculum. High inoculum (>10^6^ CFU/mouse) or a low inoculum (<10^6^ CFU/mouse) of Mm were used (n=10-28 mice/group). **E**. Time course of lesion size increases and decreases for individua mice following inoculation with 2×10^6^ CFU of *Mm*. Tail lesion size was measured (mm) every 1-2 days for each mouse for up to 5.5 weeks post infection. **F, G**. Representative images of changes in tail lesions from day 14 to day 21 in mice treated with water (**F**) or imatinib (**G**) beginning at day 14. **H**. C57BL/6J mice were infected with 2×10^6^ CFU of *Mm*. Mice were then treated with 100mg/kg/day imatinib via osmotic pump starting day -1, day 7, or day 14. Treatment was administered for 7 days before mice were sacrificed for a total infection of 6 day (n= 10 mice/group), 14 days (n= 16-26 mice/group), or 21 days (n= 11-16 mice/group). **H**. Schema of administration of imatinib or water for 6, 14 and 21 days. **I-K**. Change in lesions size during imatinib or water treatment for 6 days **(I)**, 14 days **(J)**, or 21 days **(K)**. Each data point represents one individual mouse in 2-5 experiments at each time point. Statistical tests used for comparisons in **A-D** and **I-K** was a two tailed Mann-Whitney U test, with p ≤ 0.05 judged as significant; ns, not significant at the p=0.05 level. The mean +/- SD, for each group is presented to show the variance in the data.

We next examined histopathologically the tail lesions using hematoxylin and eosin (H&E) stained tail sections of the granulomas taken at day 14 with or without imatinib treatment for 7 days (Figure 2A and B). Pathology scoring of the sections was blinded to treatment received. Scores were assigned for the number of ulcers, inflammation and necrosis in the epidermis and dermis, muscle and bone (Supplemental Figure 1A-N). We did not observe significant differences between the two groups with respect to the inflammation and necrosis in epidermis and dermis (Figure 2C), muscle (Figure 2D) or bone (Figure 2E). However, the percentage of tissue displaying inflammation was significantly reduced in the mice receiving imatinib (Figures 2G-I). Acid fast staining organisms were seen within the necrotic regions of lesions regardless of treatment (Supplemental Figure 1O). Together, these data suggest imatinib does not induce changes in microscopic composition of granulomatous lesions but does reduce the amount of inflammation.

**Figure 2.**
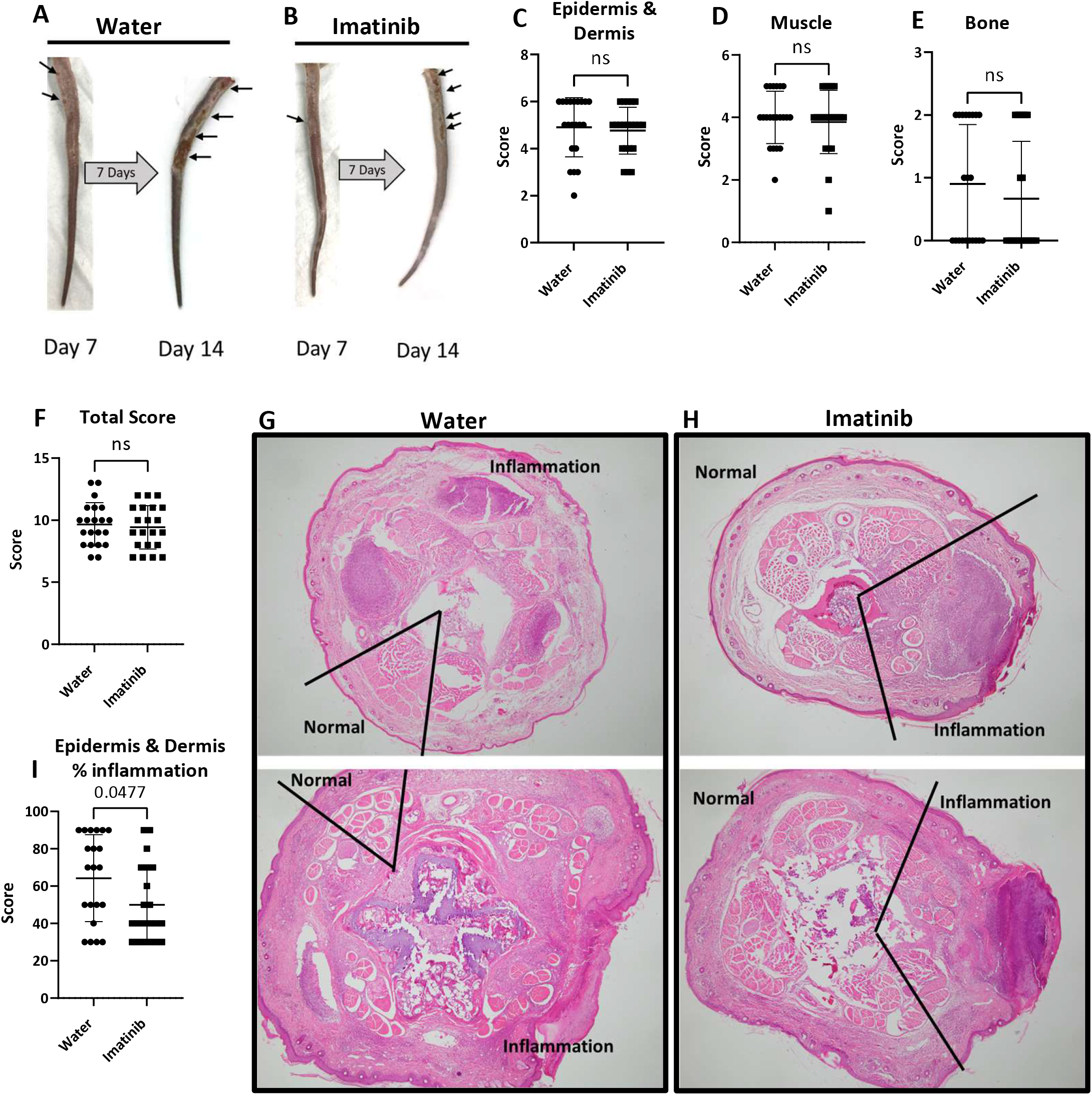
Imatinib reduces area of inflammation. Representative images of changes in tail lesions from day 7 to day 14 in mice treated with water (**A**) or imatinib (100mg/kg/day; **B**) beginning at day 7. Following sacrifice on day 14, cross sections of the tail were cut and stained with H&E. C. Composite pathology scores for the epidermis and dermis accounting for ulceration, necrosis, calcification, edema, acute inflammation, and chronic inflammation. **D**. Composite pathology scores for the muscle accounting for necrosis, edema, thrombi, acute inflammation, and chronic inflammation. E. Composite pathology scores for the bone accounting for necrosis and inflammation. **F**. Total composite pathology scores based on scores from epidermis and dermis, muscle, and bone. G,H. Representative images of whole tail sections at 14 days post infection from mice treated with water **(G)** or imatinib (100 mg/kg/day; **H**) demonstrating percent of the tissue section showing inflammation. Magnification: 40X. I. Quantification of the area of the tissue sections showing inflammation. Scoring in **C-F** and I is based on mice from 5 separate experiments (n=21 in each treatment group). Comparisons analyzed by a two tailed Mann-Whitney U test with p values ≤ 0.05 judged as significant; ns, not significant. The mean +/- SD, for each group was graphed to show the variance in the data.

### Imatinib upregulates immune genes and downregulates ECM genes in uninfected animals

To determine how imatinib affects gene expressions profiles within granulomas, RNA-seq was performed on tail sections from uninfected animals or from those infected with Mm for one or three weeks (accession no. GSE215176). Infected groups included mice treated with water or imatinib beginning one day prior to Mm infection for 7 days [“1 week infection” group (Figure 3A)] and mice in which lesions were allowed to develop for 14 days and then treated with water or imatinib for 7 days [“3-week infection” group (Figure 3A)]. RNA-seq profiles from tails of uninfected mice treated with water or imatinib served as a baseline with which to determine differentially expressed genes associated with infection based on a false discovery rate [FDR] of 0.05.

**Figure 3.**
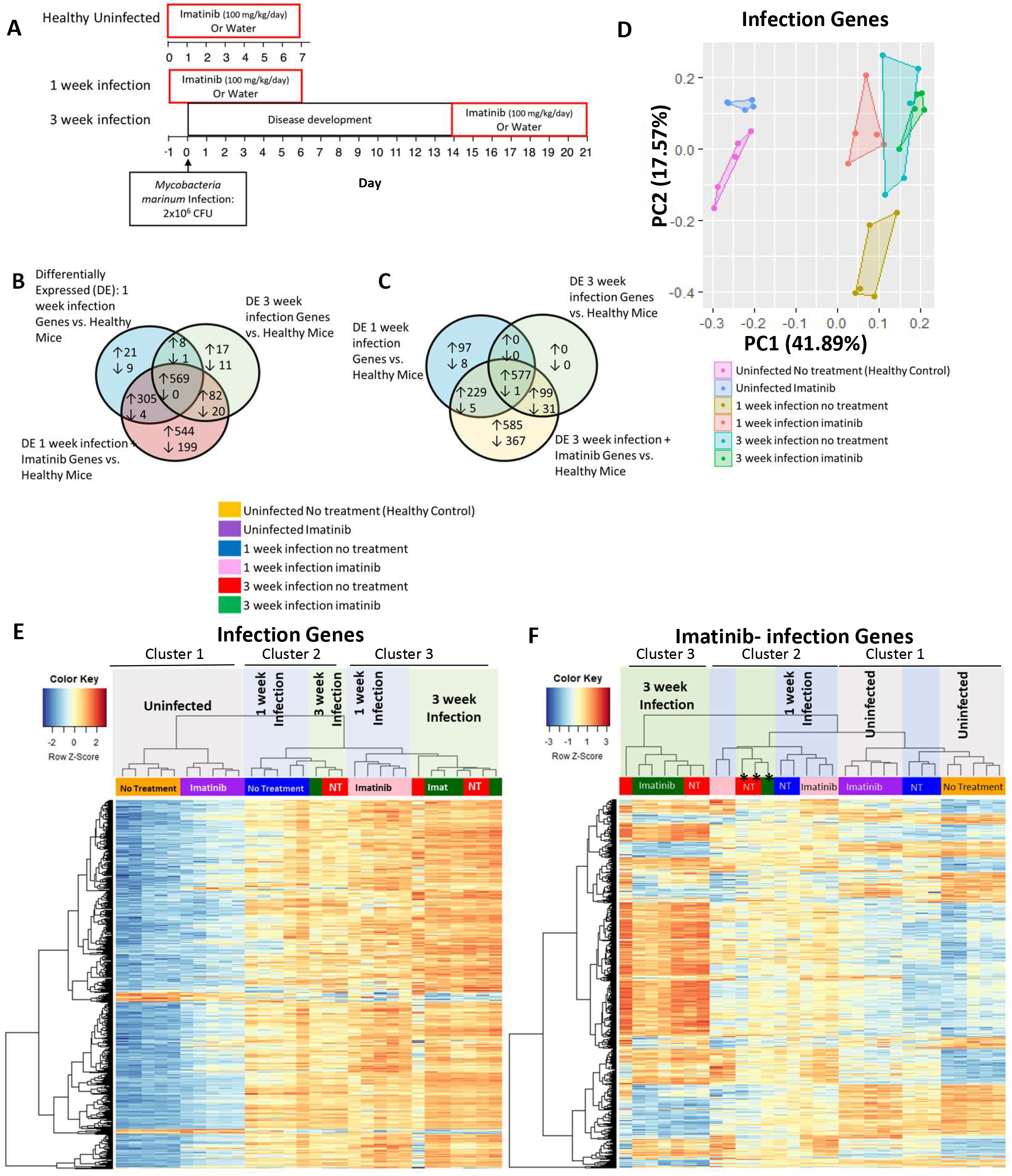
Imatinib upregulates a similar but larger number of genes with infection and at an earlier time point and with greater reliability than infection alone. **A**. Schema of administration of imatinib or water to C57BL/6J mice left uninfected or infected with 2×10^6^ CFU of *Mm*. Mice were treated with imatinib (100mg/kg/day) or water for 7 days starting day -1 or day 14 in infected group, or starting at day 0 in uninfected group, and then sacrificed seven days later (n=5 mice/group). **B. “**Infection genes” were identified by determining differentially expressed genes between the uninfected water-treated group, the one week infection water-treated group (917 genes, False discovery rate [FDR]< 0.05; blue circle), and the 3 week infection water-treated group (708 genes, FDR < 0.05; green circle) for a total of 1047 unique “infection genes” (578 genes were present at both time points). Imatinib upregulates most infection genes. Here, genes induced by imatinib at 1 week infection were identified by determining differentially expressed genes between the uninfected water treated group and 1 week infection plus imatinib group (1723 genes, FDR < 0.05; red circle). **C**. Genes induced by imatinib at 3 weeks post infection were identified by determining differentially expressed genes between the uninfected water-treated group to 3 week infection plus imatinib group (1894 genes, FDR < 0.05; yellow circle). This group was then compared to Infection genes at 1 week (blue circle) and 3 weeks (green circle) previously defined in **B. D**. The top two principal components for the variance of expression levels in all 6 of our treatment/infection groups of the 1047 “Infection Genes” identified in **B** are presented. **E**. Infection genes identified in **B** were graphed in a heat map with unsupervised hierarchical clustering. **F**. Genes specific to imatinib treatment at both time points identified in **B** and **C**, defined as “Imatinib-infection Genes,” were graphed in a heat map with unsupervised hierarchical clustering.

In the absence of infection, imatinib differentially regulated 514 genes compared to water controls, of which 373 were upregulated and 141 were downregulated. GO analysis of upregulated genes indicated that imatinib regulated diverse cellular processes, many of which were associated with the function, activation, or regulation of the immune response (Supplemental Figure 2A). Imatinib has been reported to induce myelopoiesis in mice at the dose used(46) ; accordingly, GO terms associated with neutrophil function (granulocyte activation and migration) and myeloid cell differentiation were upregulated with imatinib. GO processes significantly downregulated with imatinib include skin-epidermis development and ECM organization, which was ascribed to a significant reduction of 17 collagen subunits including *col3a1*, c*ol1a2, col1a1, col5a1*, and *col8a2* (Supplemental Figure 2B). Thus, without infection, imatinib upregulated genes and processes associated with immune function, and downregulated genes associated with ECM.

Imatinib rapidly induces expression of infection genes. Genes regulated by infection (“infection genes”) were identified as those differentially expressed at one week or three week time points following infection as compared to expression levels in uninfected mice treated with water (controls). We identified 903 genes upregulated at the one-week time-point, of which 577 remained upregulated at three weeks (Figures 3B & C). GO analysis of the infection genes upregulated at one week indicated processes involved in immune responses and cytokine production, including responses to interferon-gamma and positive regulation of TNF, IL6, and IL1β production (Supplemental Figure 2C). At the three-week time point, 99 additional upregulated genes were identified as infection genes, for a total of 676 genes at this time point. GO analysis of upregulated infection genes at three weeks indicated immune-related processes, including regulation of the cytokines IL6, IL1, IL12, IL10, IL8 and TNF, as well as macrophage activation, and responses to wounding (Supplemental Figure 2D). GO analysis of genes downregulated by infection at either the one-or three-week timepoints did not identify any processes.

We next examined how imatinib impacted expression of infection genes. To do this, the variance of infection genes without or with imatinib was analyzed by principal component analysis ([PCA]; Figure 3D). Uninfected mice clustered together in terms of PC1 (pink box), and imatinib treatment for one week shifted the variance, though only in PC2 (compare pink and blue boxes; Figure 3D). Infection alone for one week shifted the variance primarily along the PC1 axis (gold box; Figure 3D), and imatinib treatment again shifted the variance only in PC2 (compare gold and red boxes; Figure 3D). Infection for three weeks shifted the variance in PC2 (compare gold and teal boxes; Figure 3D) to a position near that seen with imatinib treatment at one week (compare red and teal boxes; Figure 3D). Imatinib treatment at this time point produced little additional shift (compare green and teal boxes), however, less variance in expression was apparent, as indicated by the reduced area of the green box. Overall, imatinib altered the pattern of variance in infection genes such that the variance at the one-week time-point with imatinib resembled that seen at three weeks without imatinib, and at three weeks, imatinib further restricted the variance.

A similar pattern was evident when expression levels for the infection genes were displayed on a heatmap with unsupervised hierarchical clustering (Figure 3E). Three main clusters were evident. Uninfected mice displayed low expression levels for most infection genes, and imatinib induced a marginal increase in expression of some of these genes (Cluster I). A second cluster included the one-week infection water-treated group together with a few mice from the three-week infection time-point, which displayed a low bacterial burden (Cluster 2). A third cluster included all the one-week infection imatinib-treated mice, together with the remaining three-week infection mice (Cluster 3). As in the PCA plot, four of the five imatinib-treated mice in the three-week infection group displayed similar gene expression profiles, whereas expression levels of infection genes in water-treated mice exhibited more variance between animals. Taken together, these data indicate that treatment with imatinib caused the pattern of gene expression at 1-week of infection to resemble that seen at three weeks, and imatinib at three weeks of infection resulted in more uniform expression of infection genes.

### Imatinib augments the infection gene signature

We next identified genes differentially regulated by imatinib. When directly comparing differences in gene expression with infection and with or without imatinib at the one- and three-week timepoints, no genes were identified as differentially expressed with imatinib at either timepoint as defined by an FDR of less than 0.05. Thus, treatment with imatinib did not cause significant changes in genes expressed during infection.

We next compared genes expressed with imatinib and infection at the one- and three-week time points to that of water-treated uninfected controls [called imatinib genes]. At one-week infection time point, imatinib upregulates 1,418 genes compared to 903 infection genes in infected mice treated with water alone (compare blue and red circles in Figure 3B). Of the 1,418 genes, 874 were shared amongst the infection and infection plus imatinib groups, and 569 were core infection genes that were upregulated at both one and three weeks of infection (Figure 3B). At the three-week timepoint, imatinib upregulated 1,490 genes, including all 676 three-week infection genes upregulated with water, as well as 229 additional genes also upregulated at the one-week time point (Figure 3C). Thus, at each timepoint, most infection genes are induced by imatinib, together with an additional 544 genes at one week and 585 genes at three weeks. Of the additional genes, ∼300 genes are specific to each timepoint, and 227 genes are upregulated by imatinib at both timepoints (Supplemental Figure 2E). Thus, imatinib induced some 902 additional genes in infected animals, not originally identified as infection genes (“imatinib-infection genes”).

When expression of the imatinib-infection genes differently regulated with imatinib during infection but not with infection alone are displayed in a heat map with unsupervised hierarchical clustering, three clusters were evident (Figure 3F). Cluster 1 contained all the uninfected mice together with 3 of 5 one-week infection mice with no treatment. The second cluster (Cluster 2) contained the rest of the mice infected for one week, including all the one-week infection mice treated with imatinib, and a few of the three-week infection mice that had lower bacterial CFUs in the spleen and tail (*, Figure 3F). The third cluster contained all remaining three-week infection mice, including those treated with imatinib or water. Taken together, these data indicate that imatinib treatment augments the infection gene signature to include additional imatinib-infection genes at both the one-week and three-week time points, which are similarly regulated without imatinib treatment, but not as reliably expressed.

We surmised that the imatinib-infection gene signature might correlate with changes in granuloma formation. To test this possibility, we further characterized the imatinib-infection gene signature, focusing on the one-week time point, where differences between the imatinib- and water-treated animals were most apparent. At this timepoint, imatinib upregulated 626 more genes than infection alone and downregulated 219 genes (Figure 4A). GO analysis of the upregulated genes indicated many immune system processes, including cell activation, response to and regulation of cytokines (Il1b, Il12, Il6, TNFα, Il8), wound healing, and cell death (Figure 4B). Notably, imatinib downregulated processes similar to those downregulated by imatinib in uninfected mice (Supplemental Figure 2B), including skin development as well as ECM and collagen organization (Figure 4C). At the three-week infection timepoint, imatinib upregulated 814 more genes than infection alone and downregulated 372 genes (Supplemental Figure 2F). GO analysis indicated the upregulated genes at three weeks are involved in similar processes as those upregulated with imatinib at the one-week timepoint. This included response to lipopolysaccharide, regulation of TNFα production, and apoptotic processes (Supplemental Figure 2G). The downregulated genes at the three-week timepoint were associated with lipid metabolism, tissue development, and supramolecular fiber organization (Supplemental Figure 2H). Thus, processes up- and down-regulated with imatinib at three weeks were similar to those evident at the one-week timepoint. Taken together, GO analysis indicated that imatinib upregulated processes associated with immune responses and wound healing, but downregulated those associated with development of epidermis and connective tissue.

**Figure 4.**
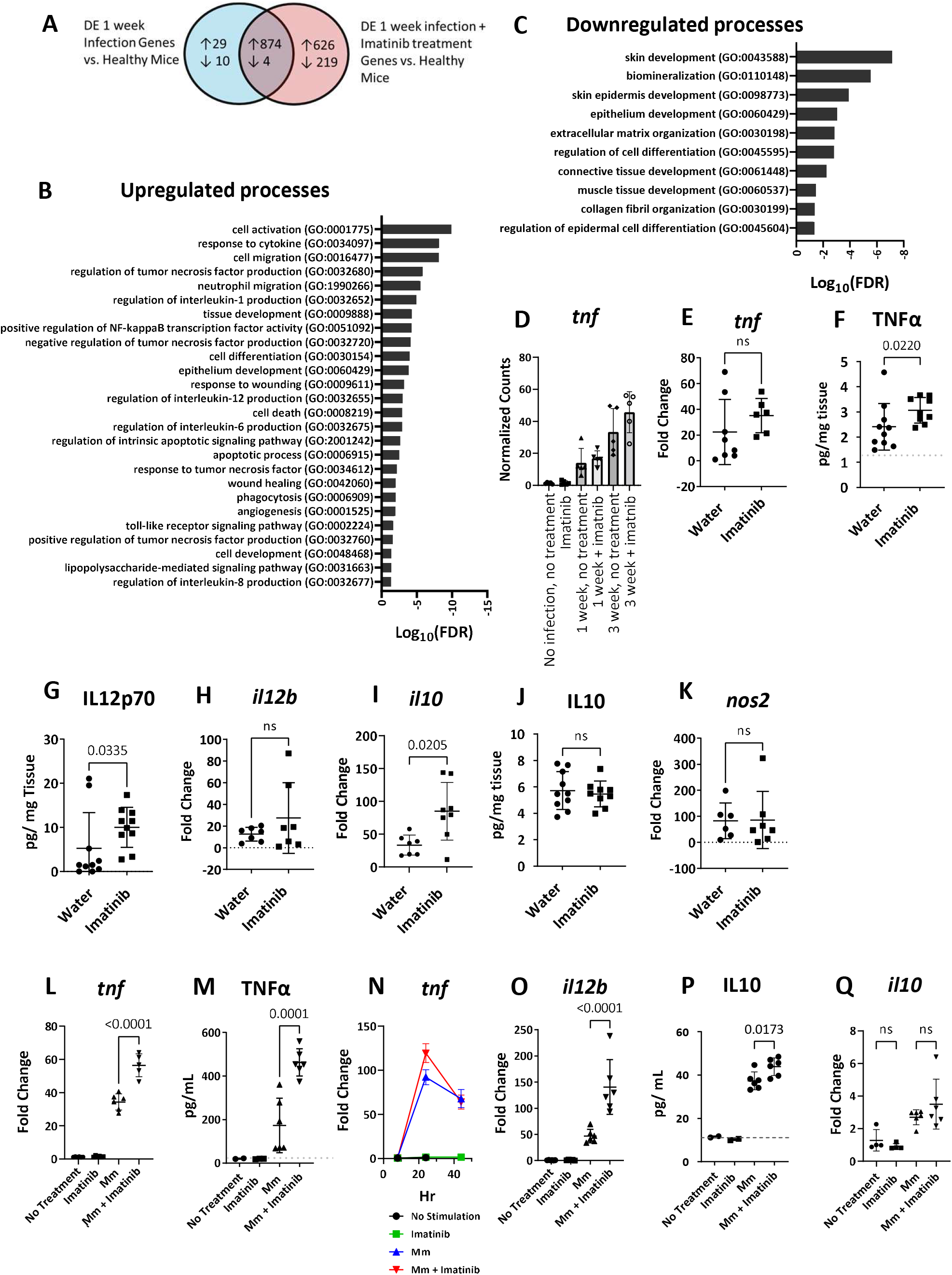
Imatinib treatment increases cell activation and cytokine production. **A**. Comparison of 1 week infection related genes and 1 week infection plus imatinib genes as described in Figure 3B. **B**. Selection of GO terms identified by analysis of 626 upregulated genes from **A** unique to imatinib treatment at 1 week infection. **C**. Selection of GO terms identified by analysis of 219 downregulated genes from **A** unique to imatinib treatment at 1 week infection. **D**. Normalized gene counts from RNAseq mapped to the *tnf* gene. **E-K**. RNA and protein were isolated from sections of the mouse tail at the 1 week infection time point (6 days infection) with water or imatinib treatment (100 mg/kg/day). qPCR was used to measure mRNA levels of *tnf* **(E)**, *il12b* **(H)**, *il10* **(I)**, and *nos2***(K)** (n=7 mice/group). Cytokine content was measured via ELISA for TNFα **(F)**, Il12p70 **(G)**, and Il10 **(J)** on the protein isolated from the mouse tails (n=10 mice/group). Statistical test used in **E-K** was a two-tailed Mann-Whitney U test, with p values indicated. A value of ≤ 0.05 was considered significant. **L-Q**. BMDMs derived from C57BL/6J mice infected with Mm at an MOI of 10 for 8, 24 or 44 hrs at 37⍰C. ELISAs were used to measure TNFα **(M)**, and Il10 **(P)** in the media at 24 hrs (n=2-6 wells/group). RNA was isolated from the cells and qPCR was used to measure mRNA levels of *tnf* **(L)**, *il12b* **(O)**, and *il10* **(Q)** at 24 hrs (n=5-6wells/ group). qPCR was used to measure levels of *tnf* in the cells 8,24, and 44hrs (**N**; n= 2 wells/ group; data are representative of 2 separate experiments). Statistical test used **in L-Q** was one-way ANOVA, with p values indicated. The mean +/- SD, for each group was graphed to show variance of data.

Imatinib treatment enhances macrophage activation and induces production of TNFα. The observations that imatinib enhances myelopoiesis and phagolysosomal fusion in infected macrophages (44, 46), and upregulates GO processes associated with immune activation (Figure 4B), led us to hypothesize that imatinib facilitates the capacity of macrophages to detect and respond to mycobacterial infection. To test this possibility, we quantified cytokine expression in granulomas. In the RNAseq analysis, infection increased levels of *tnf* at the one-week timepoint, an effect slightly augmented with imatinib, and levels increased further by three weeks, where imatinib effects were still more apparent, though these differences were not statistically significant (Figure 4D). Levels of *tnf* RNA by qPCR were likewise not significantly different with imatinib plus infection compared to infection alone at one week (Figure 4E); however, levels of TNFα protein were significantly higher with imatinib plus infection compared to infection alone (Figure 4F). Changes in the levels of *il12b* RNA, which encodes a subunit of IL12p70, were similar in both groups (Figure 4G). However, protein levels of IL12p70 were strongly upregulated with imatinib at one week of infection (Figure 4H). Levels of *il10*, which encodes the immune regulatory cytokine IL-10, significantly increased with infection plus imatinib (Figure 4I) compared to infection alone, though this effect was not recapitulated in measurements of IL10 protein levels (Figure 4J). Other factors upregulated in Mtb granulomas and associated with protection, such as *nos2*, were unchanged with imatinib (Figure 4K). Together, these data indicate that imatinib treatment activates pro-inflammatory signaling in granulomas, but also induces transcription of cytokines that resolve inflammation.

Macrophages are among the first cells infected by mycobacteria and a source of TNFα (54, 55). To test the hypothesis that imatinib induces TNFα or other cytokines in macrophages, bone marrow derived macrophages (BMDM) from WT C57Bl/6J mice were infected with Mm at an MOI of 10, exposed to imatinib, and cytokine levels in the media measured 24 hours later. Compared to untreated cells, imatinib induced expression of *tnf* mRNA (Figure 4L) and enhanced release of TNFα into the media (Figure 4M) compared to infection alone. Notably, *tnf* mRNA levels evident with imatinib returned to baseline levels similar to those of control infected cells by 44 hours p.i.(Figure 4N). Il12p70 protein was not produced in measurable levels from the BMDMs after Mm infection though levels of *il12b* mRNA were increased with imatinib treatment (Figure 4O). Imatinib also induced secretion of IL10 from BMDMs (Figure 4P), contrary to *in vivo* results, though *il10* mRNA levels remained unchanged in these cells (Figure 4Q). *nos2* mRNA levels were strongly induced with infection in BMDMs over the first 24 hours and continued to increase thereafter, with imatinib limiting *nos2* mRNA production (Supplemental Figure 3A and B). Despite imatinib reducing levels of *nos2*, a marker of M1 macrophage activation, M2 macrophage activation markers *chil3l3* (YM1) and *arg1* were not changed (Supplemental Figure 3C and D). These data indicate that upon infection of BMDMs, imatinib augments production of TNFα, a marker of macrophage activation, mirroring effects seen in infected tissues in the mice (Figure 4F), though *in vivo* regulation by imatinib of other factors, such as *nos2* and *il10*, was not recapitulated in BMDMs.

Imatinib increases cellular survival and reduces necrosis in infected BMDMs. TNFα signaling coordinates cell death and survival (56, 57), and regulates Mtb infections and granulomas, including in humans (27-30). The observation that imatinib augments TNFα production in infected cells raised the possibility that the drug might also regulate cell death or cell survival within lesions. To determine whether imatinib regulates apoptosis in granulomas, the level of cleaved caspase 3 was quantified by western analysis in the tails of the mice infected with Mm and treated for 7 days with water or imatinib.

Levels of cleaved caspase 3 were evident with infection, with some animals showing higher levels than others; however, on average, imatinib treatment did not affect levels of cleaved caspase 3 (Figure 5A), nor, by inference, the level of apoptosis. Although necrosis was evident in most tissue sections (Supplemental Figure 1A), no differences were evident with imatinib treatment (Supplemental Figure 1C, H, and M).

**Figure 5.**
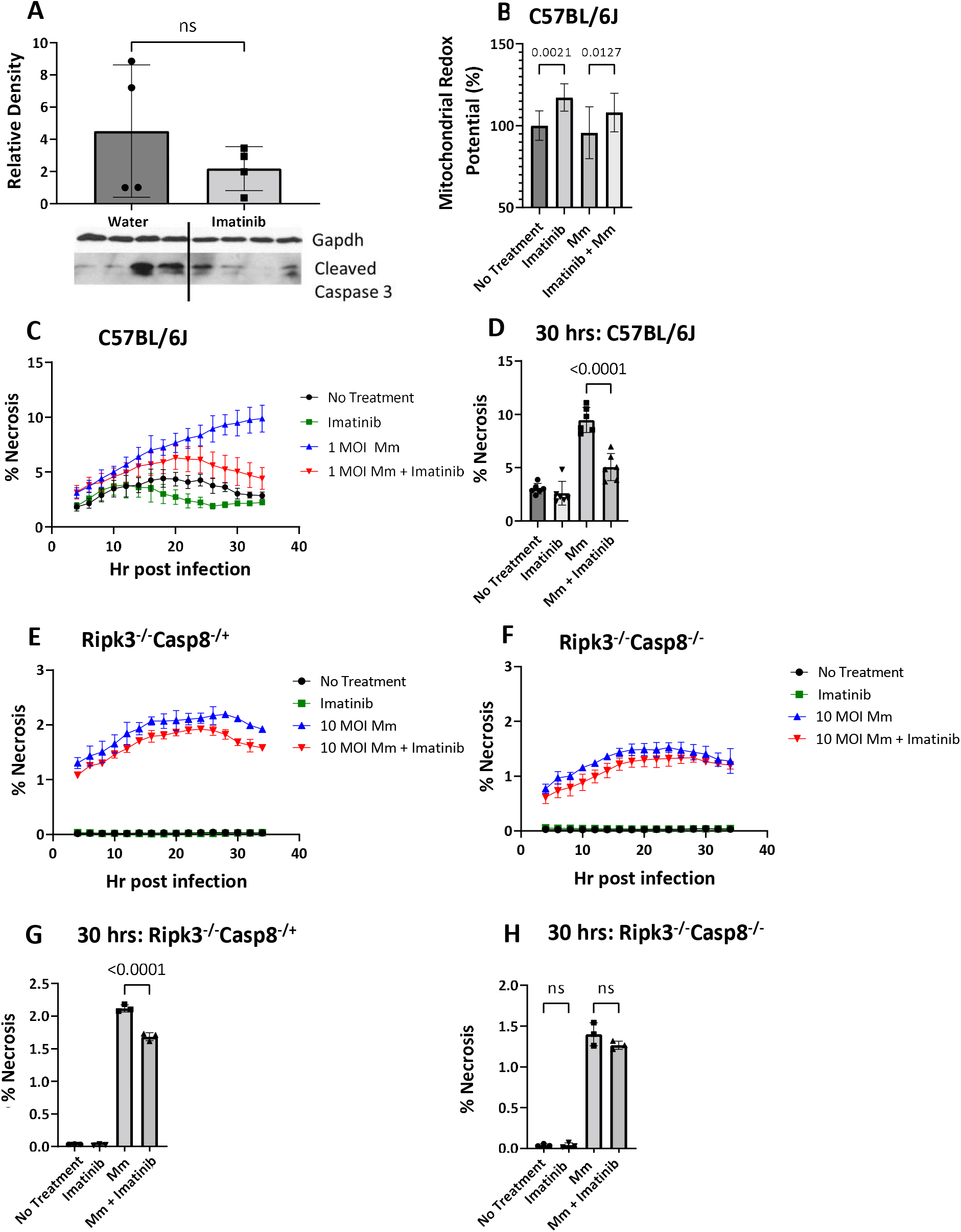
Imatinib limits necrosis in BMDMs, which depends upon caspase 8. **A**. Western blot analysis of cleaved caspase 3 was performed on protein isolated from sections of the mouse tail at 6 days post infection with water or imatinib treatment (100 mg/kg/day) for 7 days. **B**. BMDMs derived from C57BL/6J mice were infected with Mm at an MOI of 5 and left untreated or treated with 1μM imatinib. Cell viability was assessed 20 hrs post infection by measuring fluoresecence of resorufin, and indicator of redox potential. All groups were normalized to the average of the no infection/no treatment group (n= 10 wells/group; data shown are representative of 3 separate experiments). **C**,**D**. BMDMs derived from C57BL/6J mice were infected with Mm at an MOI of 1 and left untreated or treated with 1μM imatinib. Cells were imaged every 2 hrs with Cytotox dye (n= 6 wells/group, representative of 3 separate experiments), an indicator of necrosis. Percent (%) necrosis was quantified at each time point and graphed as a function of time post infection (**C**). Statistical analysis was performed on data at 30 hrs post infection, and graphed in **D. E-H**. BMDMs derived from Ripk3^-/-^ Casp8^+/-^ or Ripk3 ^-/-^ Casp8^-/-^ mice were infected with Mm at an MOI of 10 and left untreated or treated with 1μM imatinib. Cells were imaged every 2 hrs with Cytotox dye (n=3 wells/ group, representative of 3 separate experiments). Data is show as a function of time **(E**,**F)**, and at the 30 hr timepoint **(G**,**H)** for the Ripk3^-/-^ Casp8^+/-^ or Ripk3 ^-/-^ Casp8^-/-^ BMDMs, respectively. Statistical test used in **A** was two-tailed Mann-Whitney U test. Statistical test used in **B, D, G, & H** was one-way ANOVA. p values are indicated in graphs. The mean +/- SD for each group was graphed to indicate variance of data.

Induction of NFKb signaling by TNFα upregulates expression of factors that promote cell survival (57, 58), raising the possibility that imatinib might promote cell survival. To test whether imatinib increased cell viability, mitochondrial redox potential was assessed in WT BMDMs. To do this, cells were treated with imatinib with or without infection with Mm at an MOI of 5, and mitochondrial redox potential measured 24 hours later. Imatinib treatment of uninfected BMDMs increased redox potential by ∼15%, and no changes in were evident upon infection with Mm compared to uninfected cells. However, imatinib treatment increased redox potential by 8% in infected cells (Figure 5B), indicating a survival benefit with the drug.

To determine whether imatinib affected necrosis, WT BMDMs were cultured with or without imatinib and infected with Mm at an MOI of 1 or left uninfected. Cells were imaged over a 36-hour period in the presence of Cytotox reporter, which labels cells that have lost membrane integrity, an indication of necrosis. With or without imatinib, some necrotic cell death was evident after four hours p.i., which plateaued after 24 hours. Imatinib reduced the amount of necrosis between 16 and 24 hours (compare black and green lines; Figure 5C). Upon infection with Mm at an MOI of 1, the percentage of cells undergoing necrosis increased over time (blue line, Figure 5C). With imatinib, the levels of necrosis in infected cells initially increased over 20 hours at a rate similar to that of untreated cells. After 24 hours, levels of necrosis in untreated cells continued to increase whereas necrosis in imatinib-treated cells declined to levels seen in uninfected cells (compare red and blue lines; Figure 5C). By 30 hours p.i., the level of necrosis with infection was 2-fold lower in imatinib-treated cells compared to controls (Figure 5D). Taken together, these data indicate that imatinib increases survival of BMDMs and limits necrosis.

Imatinib effects on necrotic cell death in BMDMs depend upon caspase 8. To test the possibility that imatinib might regulate cell death or survival, imatinib effects were assessed in BMDMs derived from mice with deficiencies in TNFα signaling. Cells from mice lacking RIP3 kinase but heterozygous for caspase 8 (Ripk3^-/-^ Casp8^-/+^) do not undergo TNFα-mediated necroptosis, whereas cells from mice lacking both RIP3 kinase and caspase 8 (Ripk3^-/-^ Casp8^-/-^) can neither undergo necroptosis nor extrinsic apoptosis in response to TNFα (59, 60). Without infection, little necrotic cell death was evident in BMDMs derived from either Ripk3^-/-^ Casp8^-/+^ mice or Ripk3^-/-^ Casp8^-/-^ mice, and no additional effect of imatinib was discernable (Figures 5E and F; Black and green lines). Induction of discernable levels of cell death in BMDMs from Ripk3^-/-^ Casp8^-/+^ or Ripk3^-/-^ Casp8^-/-^ required infection with Mm at an MOI of 10 rather than an MOI of 1 used in WT BMDMs, though even at this MOI, the percentage of cells undergoing necrosis was still lower than in WT cells (compare Figure 5C, E, and F). With infection, BMDMs from Ripk3^-/-^ Casp8^-/+^ and Ripk3^-/-^ Casp8^-/-^ showed similar initial levels of necrosis without imatinib, which increased over 24hrs before plateauing (Figures 5E and F). With imatinib treatment, Ripk3^-/-^ Casp8^-/+^ showed statistically significant reductions in the amount of necrosis by 30 hrs. p.i. (Figures 5E and G; compare blue and red line). However, imatinib did not affect the level of necrosis in Ripk3^-/-^ Casp8^-/-^ cells (Figure 5F and H, compare blue and red lines). Together, these data indicate that infection with Mm induces necrosis in BMDMs, and that imatinib limits necrosis in a manner that depends upon caspase 8.

Imatinib-mediated reduction in pathology of granulomatous lesions is abrogated in mice lacking caspase 8. We next determined effects of imatinib on granuloma growth in Ripk3^-/-^ Casp8^-/+^ and Ripk3^-/-^ Casp8^-/-^ mice infected with Mm and exposed to imatinib or water seven days later (Figure 6A). Lesions in Ripk3^-/-^ Casp8^-/+^ mice treated with water grew an average of 13.5mm from day seven to fourteen p.i, whereas lesions from mice treated with imatinib grew on average by 4mm over the same period (Figure 6B). Lesion growth in Ripk3^-/-^ Casp8^-/+^ mice was not significantly different from that seen in C57Bl/6J mice, and imatinib treatment reduced lesion growth by a similar amount in both the C57Bl/6J and Ripk3^- /-^ Casp8^-/+^ animals (Figures 6B and C). Lesions in infected Ripk3^-/-^ Casp8^-/-^ mice grew by 23mm on average from day seven and fourteen, with half the animals developing severe and extensive lesions measuring >30mm. Treatment of the Ripk3^-/-^ Casp8^-/-^ mice with imatinib did not significantly reduce lesion growth (Figures 6B and D). The bacterial burden in the tails of both Ripk3^-/-^ Casp8^-/+^ and the Ripk3^-/-^ Casp8^-/-^ mice was ∼7 Log _10_ CFU gram^-1^tissue, and imatinib treatment resulted in a 1 log_10_ (Figure 6E) increase in both genotypes. In the Ripk3^-/-^ Casp8^-/+^ mice, this increase in CFUs in the tail was seen despite the reduction in lesion size. Notably, bacteria load in spleen was similar between C57BL/6J, Ripk3^-/-^ Casp8^-/+^, and the Ripk3^-/-^ Casp8^-/-^ mice, and imatinib did not cause a detectable change (Figure 6F). These data indicate that deletion of RIP3 kinase does not preclude imatinib from limiting the pathology associated with granulomatous lesions; moreover, the capacity of imatinib to limit pathology of granulomatous lesions resulting from Mm infection was abrogated in mice lacking caspase 8.

**Figure 6.**
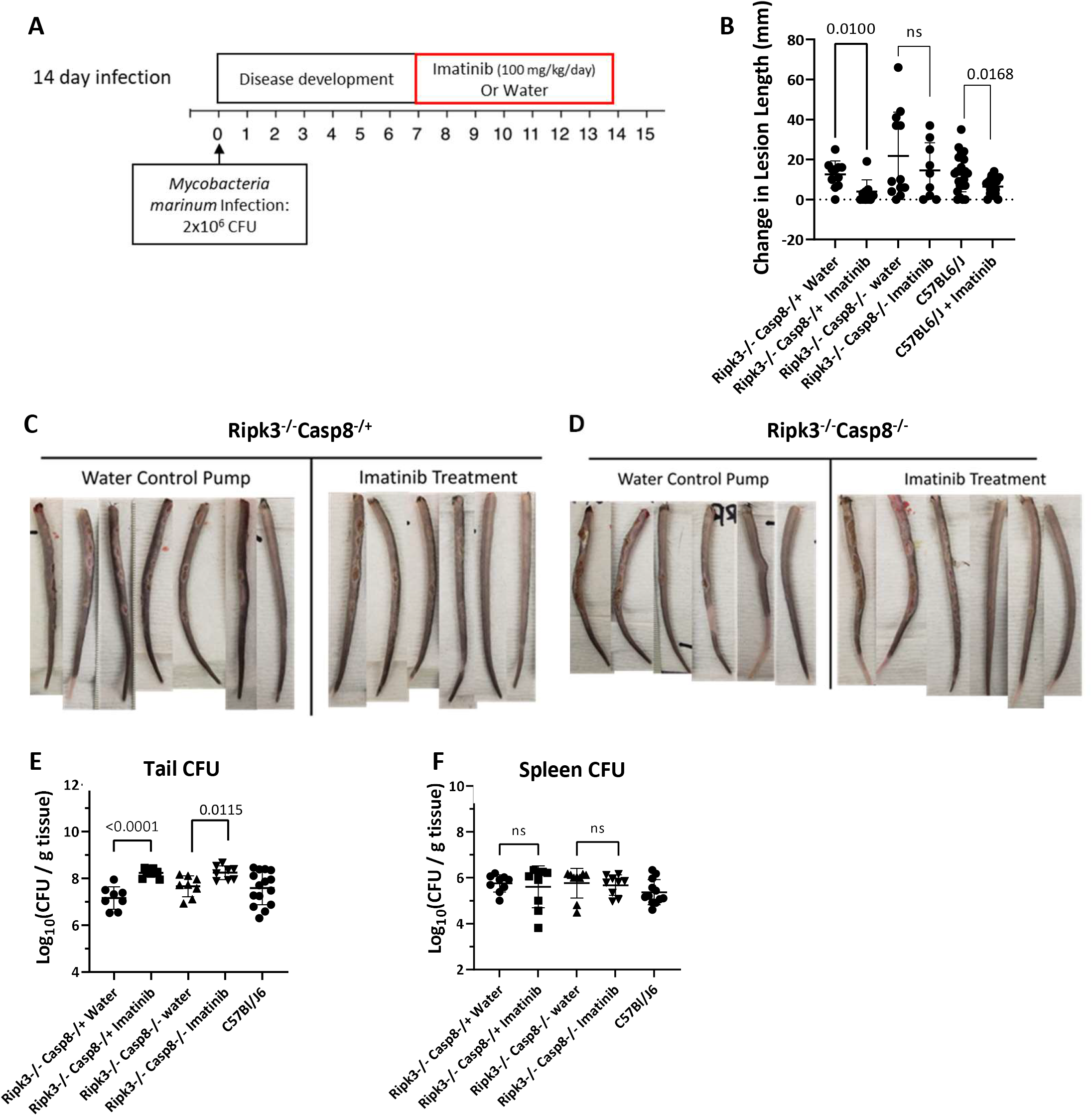
Imatinib-mediated reduction in pathology of granulomatous lesions is abrogated in mice lacking caspase 8. **A**. Ripk3^-/-^ Casp8^-/-^ mice or Ripk3^-/-^ Casp8^-/+^ mice were infected with 2×10^6^ CFU of *Mm*, for 7 days. Mice were then treated with imatinib (100mg/kg/day) or water starting day 7 post infection. for an additional 7 days before mice were sacrificed at day 14 (n= 9-12 mice/group). **B**. Lesions were measured at the beginning of treatment and at the end of treatment to determine the change in lesion size over the treatment period. Historical data from Mm-infected C57/Bl6/J mice treated with water or imatinib (100 mg/kg/day) was included for comparison. **C**,**D**. Representative images of the tail lesions from Ripk3^-/-^ Casp8^-/+^ **(C)** and Ripk3^-/-^ Casp8^-/-^ **(D)** mice. **E**. Tail CFUs from the Ripk3^-/-^ Casp8^-/+^ and Ripk3^-/-^ Casp8^-/-^ treated with water or imatinib and historical control from C57BL/6J mice infected as above. **F**. Spleen CFUs from the Ripk3^-/-^ Casp8^-/+^ and Ripk3^-/-^ Casp8^-/-^ with no treatment or with imatinib treatment and historical control from C57BL/6J mice infected as above. Statistical test used in **B**,**E**, and **F** was two-tailed Mann-Whitney U test, with p values indicated. The mean +/- SD, for each group was graphed to show variance of data.

## Discussion

Imatinib augments anti-mycobacterial immune responses. Imatinib both induces myelopoiesis and promotes phagolysosome fusion (44, 46). Data presented here indicate that many untreated mice develop transcriptional responses associated with immune activation after three weeks of infection that resemble those induced in infected animals after one week with imatinib (Figure 3). Moreover, imatinib-induced transcriptional responses at three weeks appear more uniform owing to an increased proportion of mice displaying similar responses at that time point. In short, the response to infection with imatinib is more rapid and more efficient, but not substantially different from that developing at three weeks without drug. One possible explanation for the more rapid response to infection with imatinib is an increased capacity for phagolysosomal fusion in infected macrophages (44, 45), which precludes the capacity of mycobacteria to limit macrophage activation and antigen presentation. Additionally, increased numbers of innate immune cells resulting from enhanced myelopoiesis with imatinib could facilitate a more robust innate response (46).

TNFα appears to be a key marker of macrophage activation in response to imatinib (Figure 4). However, granuloma formation and maintenance are also in part mediated by TNFα. Thus, mice deficient in TNFα exhibit disorganized granulomas (27, 28), and patients administered drugs that neutralize or inhibit TNFα are at increased risk of reactivating TB (29-31). Other cytokines, including interleukin-12 (Il12) and interferon-γ (INF-γ), also contribute to development and organization of granulomas, and activate innate immune cells that combat Mtb (33, 55, 61-63). In line with TNFα as a cellular activation marker, imatinib initially increases TNFα production in infected BMDMs. However, induction of TNFα is not sustained, and declines to levels seen in untreated cells (Figure 4). Notably, sustained increases in TNFα are associated with chronic inflammation and tissue damage in inflammatory bowel disease and rheumatoid arthritis (57). Likewise, excess TNFα is associated with ROS-mediated tissue damage in Mm infections in zebrafish (64, 65). Taken together, these data support the hypothesis that imatinib subtly modulates TNFα production to augment anti-mycobacterial immune responses but may also regulate granuloma growth.

Imatinib limits granuloma formation and growth. Granulomas have long been considered a beneficial response against mycobacterial infection as they limit bacterial dissemination. However, during Mtb infection, these structures have also been shown to promote chronic infection by limiting access of immune cells to bacteria(66, 67). Granulomas also limit access of antibiotics to the bacteria, resulting in suboptimal concentrations in the caseum, where most bacteria reside (37, 38). Bacterial persistence can also induce chronic inflammation, which can scar and damage lung tissue. Moreover, attempts to heal damaged tissue results in deposition of collagen, which causes fibrosis and restricts pulmonary elasticity (68). In mice infected with Mm, where granulomas are induced by a high inoculum (Figure 1), significant reductions in lesion growth are evident with imatinib. This effect does not depend on bacterial load, which is unaffected by imatinib at this inoculum. Microscopic analysis indicated that imatinib reduced the amount of inflammation but not the composition as necrosis, calcification, edema, ulceration, thombi, acute inflammation, and chronic inflammation were observed in both groups (Figure 2). Taken together, these data indicate that imatinib reduces the growth of the lesions and the extent of inflammation but does not otherwise alter histological features of the lesions.

Imatinib has been shown to limit collagen deposition by inhibiting PDGFR and fibrosis in non-infectious indications (48-53). Accordingly, reductions in collagen are suggested by the GO analysis of gene expression data from imatinib-treated mice (Supplemental Figure 2). These data raise the possibility that imatinib may facilitate wound healing in a way that is less likely to induce fibrosis, which in turn may limit scaring and long-term tissue damage. This finding has important implications for lung health during Mtb infection. Chronic inflammation has been associated with increased fibrosis and lung dysfunction in TB patients (36, 39, 40, 47), which results in increased economic burden of patients who have successfully completed antibiotic therapy(42). We postulate that imatinib may be an effective treatment for patients with active TB by reducing fibrosis and lung dysfunction. The structure of the granuloma has also been shown to limit antibiotic penetrance into the granuloma(37, 38). By limiting formation and/or promoting resolution of granulomas, imatinib may increase access of antibiotics to the bacteria and thereby shorten duration of treatment.

Imatinib promotes cell survival and limits necrosis. Cell death is a major contributing factor to bacterial spread and tissue pathology (33, 69-71). Apoptosis has been shown to limit spread and replication of Mtb by encapsulating the bacteria in apoptotic bodies and allowing macrophages to efferocytose apoptotic bodies containing Mtb, and transfer them into the lysosome (72). By contrast, necrosis facilitates spread of bacteria, promotes inflammation, and contributes to tissue damage by releasing ROS (33, 71, 73). Ferroptosis, an iron-dependent form of necrosis, has recently been identified as a major contributor to pathology during Mtb infection (73). Several lines of evidence indicate that altering the type of death that a cell undergoes can impact Mtb infection. For example, a mutation in the Lpr1 gene, which encodes LDL receptor, can shift macrophages from necrosis to apoptosis, thereby limiting replication and spread of the bacteria (70). In this regard, imatinib has been shown to increase apoptotic cell death in BCR-Abl^+^ transformed cells but is generally without effect on non-transformed cells (74). Tails in mice infected with Mm display reduced gross pathology and more focused areas of tissue damage with imatinib, which is consistent with reduced necrosis and/or increased survival though not changes in necrosis at a gross level were evident (Supplemental Figure 1). Accordingly, in cultured macrophages, imatinib has little effect on uninfected cells, but reduces necrosis in infected ones (Figure 5).

Our data with the Ripk3^-/-^ Casp8^-/-^ mice suggest a possible role for caspase 8 in imatinib effects on granulomas (Figure 5 and 6), possibly by regulating necrosis, or cell survival, or both. In this regard, caspase 8 promotes extrinsic apoptosis in response to signaling via death receptors such as TNF receptor 1 and CD95 (57, 75), though no evidence of caspase 8-mediated apoptosis was evident in our analysis (Figure 5A). However, caspase 8 also promotes cell survival by complexing with c-Flip, which inhibits apoptosis, and precludes activation of RipK3, which prevents necroptosis (76, 77). Finally, caspase 8 regulates transcriptional responses that dampen inflammation (78). Any of these activities could contribute to imatinib’s effect on reducing granuloma growth or limiting inflammation *in vivo*. Further characterization of cellular mechanisms associated with enhanced survival and/or reduced necrosis with imatinib and caspase 8 are needed to elucidate how the drug facilitates resolution of granulomas.

Interestingly, an increase in CFUs was observed in the mouse tail but not spleen with imatinib treatment in both mice strains lacking RipK3. In mice infected with Mtb, it has been suggested that RipK3 facilitates the spread of bacteria (79) by mediating necrotic death, and mice lacking RipK3 developed lower bacterial CFUs in their lungs. In our model system, mice deficient in RipK3 did not show significant differences in bacterial colonization of the tails compared to control C57BL/6J mice. While imatinib treatment increased the bacteria load in the tail, the bacteria load induced by imatinib was not significantly different from that in C57BL/6J mice. Based on our analysis to date, a role for RipK3 in imatinib effects on Mm is not apparent. However, we cannot rule out that possible secondary alterations in immune responses evident in these mutant animals that could mask an addition role for this enzyme during infection.

Imatinib as an HDT. Recent efforts by us and others (80) have raised the possibility that HDTs may be effective against TB. Because HDTs do not directly select against the bacteria, they are less likely to engender resistance compared to antibiotics. Some HDTs interfere with the capacity of Mtb to subvert host systems, whereas others regulate immune responses, so as to induce novel responses or target deficiencies in individuals with active disease (81). Drug discovery efforts have traditionally focused on drugs targeting specific processes with minimal off-target effects, though recent efforts, for example with COVID-19, have targeted hyperinflammatory pathways as a means to limit tissue damage(81). Such approaches can be complex to implement, requiring proper timing and dosing to limit tissue damage while still permitting requisite inflammatory responses that contain the infection. Our studies suggest that imatinib likewise affects multiple aspects of the host immune response to mycobacteria, but it does so in ways that do not fundamentally alter the response, but rather subtly tune it to favor the host.

Imatinib remains a promising candidate HDT to treat mycobacterial infections such as TB by not only activating innate immune responses but also reducing pathology. Imatinib is currently being tested in the IMPACT-TB clinical trial to determine effective dosing in healthy humans as measured by the capacity of human blood to eliminate mycobacteria. Data presented here indicate that imatinib does not fundamentally alter the immune response to mycobacterial infection, but rather augments the rate and efficiency at which it develops. This work also shows how an HDT can limit granuloma formation. Such an effect has important implications for therapeutic strategies to control TB as the granuloma has been shown to limit antibiotic and immune cell access to the bacteria, leading to bacterial persistence and contributing to lasting tissue damage and scaring.

## Acknowledgments

The authors thank Rustom Antia, Greg Bisson, Robert Wallis, Deepak Kaushal, Ruth Napier, Alyson Swimm, Bernie Weiss, and Robert Sonowal for helpful discussions or for reviewing the manuscript. Research reported in this publication was supported in part by the Cancer Tissue and Pathology shared resource of Winship Cancer Institute of Emory University and NIH/NCI under award number P30CA138292. Research reported in this publication was supported by the Emory NPRC Genomics Core in part by NIH P51 OD011132. Sequencing data was acquired on an Illumina NovaSeq 6000 funded by NIH S10 OD026799.

## Materials and Methods

### Mice

8–12-week-old C57Bl/6J mice were purchased from The Jackson Laboratory. Ripk3^-/-^Casp8^-/-^, or Ripk3^-/-^Casp8^-/+^ were bred at Emory University by crossing Ripk3^-/-^ Casp8^-/+^ females to Ripk3^-/-^Casp8^-/-^ males (60). PCR genotyping of Casp8^−/−^ and Casp8^-/+^ mice was performed with primers 5⍰- TTGAGAACAAGACCTGGGGACTG and 5⍰-GGATGTCCAGGAAAAGATTTGTGTC. PCR amplification allele produces a 750-bp band (Casp8^+^), or a 200-bp band (Casp8^−^). Mice were bred and maintained by Emory University Division of Animal Resources where all procedures were approved by the Emory University Institutional Animal Care and Use Committee.

### Imatinib administration

Imatinib mesylate salt was dissolved in water and loaded into Alzet pumps (Braintree Scientific, 1007D; Cupertino, CA) capable of dispensing a continuous flow of drug at 100mg/kg/day. Pumps were inserted subcutaneously into anesthetized 8-12-week old mice. Alzet pumps were inserted 24 h prior to infection or 7-21 days post infection depending on experiment time course.

### Bacterial strains

M. marinum (Mm) strain 1218R (ATCC 927) was grown in Middlebrook 7H9 broth (7H9) (BBL Microbiology Systems, Cockeysville, MD) supplemented with ADC (Difco Laboratories, Detroit, MI,) and 0.05% Tween 80 (Sigma-Aldrich, St. Louis, MO) or 0.025% Tween 80. For CFU assays, 7H10 agar supplemented with 10% oleic acid-albumin-dextrose-catalase (OADC) was used (Difco Laboratories, Sparks, MD). For Mm infection in mice and infections of BMDMs, bacterial stocks were grown at 30°C for 2 days to an OD_600_ of 0.4 approximately 6.3×10^6^CFU/mL (Eppendorf, BioPhotometer; Hamburg, Germany). Bacteria were washed with sterile PBS and aspirated through a 27G needle against the side of the tube multiple times to break up bacterial clumps. Bacteria was then diluted in sterile phosphate buffered saline [PBS] for mouse infection or complete cell culture media for the BMDM infections.

### Mouse Mm infection

Mice were infected with actively growing Mm (ATCC 927) via the tail vein to induce a systemic infection that results in the development of mycobacterial granulomas on the tail of the mouse. The inoculum, determined by retrospective plating, was ∼2×10^6^CFU in each experiment. In C57BL/6J mice tail lesions appear on the mouse tail about 1 week after infection and can be monitored over time by measuring the length of each lesion from top to bottom using a ruler, to get a measure of the total lesion lengths per mouse tail. After 7 days of treatment, tail, spleen, and lung were harvested. For CFU, spleens, and lung, were weigh and homogenized in the Bullet Blender Tissue homogenizer (Next Advance, Troy, NY) in 1 ml PBS. ∼3-5 mm of tail was cut into small pieces using sterile scissors before being homogenized in the same way as lung and spleen. Each homogenate was diluted and spread on 7H10 agar plates. Colonies were scored after 7 days at 30°C. Colonies per ml were normalized to the initial weight of the tissue, to determine CFU/g tissue.

### Tail protein isolation for ELISA and Western analysis

For soluble protein isolation for ELISA, 3-5mm sections of tail were cut fresh from the tail and immediately placed in liquid nitrogen. Samples were stored at -80°C until processed. Samples were weighted and put in PBS, scissors were used to break apart the tissue before the tissue was homogenized in the Bullet Blender Tissue homogenizer (Next Advance, Troy, NY). Il12P70 (Invitrogen), TNFα(Invitrogen), and IL10 (Invitrogen) ELISAs were run according to manufacture directions. For western analysis of tail protein content, 3-5 mm of flash frozen tail samples were weighted and put in ice cold RIPA buffer supplemented with a protease inhibitor cocktail (Roche). Scissors were used to break apart the tissue before the tissue was homogenized in the Bullet Blender Tissue homogenizer (Next Advance, Troy, NY). Protein concentration was determined using BCA assay kit (ThermoFisher), before the sampled were run on 15% SDS-PAGE gel and blotted onto a PVDF membrane. Cleaved Caspase 3 (ASP175) antibody (1:1000, Cell Signaling) and GAPDH (D16H11) antibody (1:1000, Cell Signaling) were used sequentially on the same membrane to quantify and normalize protein content.

### Mouse tail RNA isolation

3-5mm sections of tail were cut fresh from the tail from areas containing granulomatous lesions and immediately placed in Trizol (Invitrogen). Tissue was cut with sterile scissors and then homogenized in the Bullet Blender Tissue homogenizer (Next Advance, Troy, NY). RNA extraction was performed in accordance with manufacture instructions.

### Histology

Tails were removed from the mouse and a clean razor used to slice tails into 3-5 mm lengths that were placed in a cassette and submerged in 10% neutral buffered formalin for 24-48 hours. Tail sections were washed and decalcified in Immunocal decalcifier (StatLab, McKinney, Tx) for 48 hrs before being moved to 70% ethanol for storage until the sections could be embedded in paraffin, and sectioned. Sections were stained by H&E (abcam) and acid-fast bacillus [AFB] stain (abcam).

### Histology scoring

Histology was obtained from the tails of mice infected for 14 days and treated with control or 100 mg/kg/day imatinib for the last 7 days of infection. Slides of H&E-stained tail sections containing 2-9 individual tail tissue cross sections/mouse were deidentified and randomized before they were submitted to a pathologist for scoring. The pathologist examined slides first to create a rubric for characteristics, then blindly scored all the sections based on the noted characteristics. The percent of inflammation was determined in the section with highest amount of inflammation.

### BMDM generation

L929 cells were maintained in DMEM (Corning) with 10% fetal bovine serum (FBS, Gibco), 100U/ml penicillin and 100 U/ml streptomycin (Invitrogen). At confluency, fresh media was added to the L929 cells and collected and sterile filtered after 3 days. L929 conditioned media was stored at -20°C until use. For BMDM culture, tibias and femurs were collected from C57BL/6J, Ripk3^-/-^ Casp8^-/-^, or Ripk3^-/-^Casp8^-/+^ mice. The ends of the bones were removed with a clean razor before the bones were placed in a sterile 0.7 ml Eppendorf tube with the bottom tip cut off inside a 1.5 mL sterile Eppendorf tube. Tubes were centrifuged at 13,000 RPM for 3 minutes to extrude the bone marrow cells. Cells were washed and filtered through a 70μM mesh filter with sterile PBS, before being plated in 10 cm petri dishes with 20% L929 conditioned media in fresh DMEM + 10% FBS + 100U/ml penicillin and 100 U/ml streptomycin. Cells were differentiated for 7-9 days before collection in ice cold PBS containing 0.5M EDTA.

### BMDM RNA and protein collection

For RNA and protein collection, BMDMs were plated at 1×10^6^ cells/ well in a 6 well plate. Cells were allowed to adhere overnight before being infected with Mm at an MOI of 10 and treated with 1μM imatinib. Infection commenced for 24 hrs, at which time media was removed and stored at -80°C for cytokine quantification. TNFα (Invitrogen), and IL10 (Invitrogen) ELISAs were performed according to manufacturer instructions on undiluted media. Cells were lysed and RNA was collected using the RNeasy Mini Kit (Qiagen), accordioning to manufacture instructions.

### BMDM cell viability assay

For cell viability assays, 10,000 cells/well were plated into a 96-well plate and allowed to adhere overnight. Cells were infected with actively growing Mm at an MOI of 5 and treated with 1 μM Imatinib in DMEM + 10% FBS and placed in an incubator at 37°C with 5% CO_2_. After 20 hours of infection, Cell Titer-Blue (Promega, Madison, WI) was added to the media and cells were incubated for 4 hrs at 37°C with 5% CO_2_ before fluorescence was measured using a plate reader.

### BMDM Time course experiments

For time course experiments, 1-3×10^5^ cells/well were plated into a 24 well plate and allowed to adhere overnight. BMDMs from C57Bl/6J mice were infected with actively growing Mm at an MOI of 1 of while BMDMs from Ripk3^-/-^Casp8^-/-^ and Ripk3^-/-^Casp8^-/+^ mice were infected with actively growing Mm at an MOI of 10 for 2 hrs in DMEM + 10% FBS. Wells were washed with PBS and 200 μg/ml amikacin in DMEM + 10% FBS was added for 2 hrs. Cells were washed again with PBS before DMEM + 10% FBS was added with 250nM Incucyte® Cytotox Green (Sartorius) +/- 1 μM Imatinib. Trays were then placed in the IncuCyte® ZOOM (Saritorius Essen) at 37°C with 5% CO2. Phase images and Green Fluorescence Images were taken at 20x every 2 hrs in. The area of phase objects and the area of green objects was quantified to determine the green area per phase area for each image, indicating the amount of necrosis indicated by the green stain per the area of cells.

### RNA-seq analysis

RNA was isolated from mouse tails from uninfected mice +/- 100mg/kg/day imatinib, infected mice at 6 days post infection +/- 100mg/kg/day imatinib, or infected mice 21 days post infection +/- 100mg/kg/day imatinib. RNA from the tails of 5 individual mice were used per group for a total of 30 samples. RNA libraries for RNA-seq were prepared using Clontech SmartER Stranded Total RNA-seq Kit-Pico Input Mammalian + rRNA depletion following manufacturer’s protocols. Sequencing was performed at Yerkes Nonhuman Primates Genomics Core, Emory University, using Illumina NovaSeq 6000. Quality control was performed on the raw reads using FastQC and the remaining analysis was done using R. Adaptors were trimmed from the ends of reads using QuasR. Hisat2 was used for alignment of the reads to the GRCm38.p6 mouse genome with 66-80% of reads mapped to a single gene. 6-25% of the mapped reads were successfully assigned to gene alignments resulting in 5.5 million-14million assigned alignments per sample. Genes with low expression levels (>20 copies per million) in at least 5 samples filtered out. Data was normalized using the Trimmed Mean of M-values [TMM] method. Differential expression analysis was performed using edgeR, genes were determined by comparison of each group to the uninfected mice not treated with imatinib. Raw reads and processed gene counts in this paper have been deposited in the Gene Expression Omnibus (GEO) database, https://www.ncbi.nlm.nih.gov/geo/ (accession no. GSE215176).

### QPCR

RNA was isolated from mouse tail or BMDMs as previously described. Reverse transcription was performed using the RevertAid First-Strand cDNA Synthesis Kit (Thermo Fisher Scientific) with the oligo(dT)18 primer. SYBR^®^ Green Supermix (Bio-Rad) according to manufacturer instructions using a MyiQ real-time PCR system (Bio-Rad). The ΔΔCT method was used to determine relative gene expression using Gapdh as internal controls. Primers used were: Gapdh-forward 5’ AGGTCGGTGTGAACGGATTTG3’, Gapdh-reverse 5’TGTAGACCATGTAGTTGAGGTCA3’, il12b-forward 5’TGGTTTGCCATCGTTTTGCTG3’, il12b-reverse 5’ACAGGTGAGGTTCACTGTTTCT3’, tnf-forward 5’CCCTCACACTCAGATCATCTTCT3’, tnf-reverse 5’ GCTACGACGTGGGCTACAG3’, il10-forward 5’GCTCTTACTGACTGGCATGAG3’, il10-reverse 5’CGCAGCTCTAGGAGCATGTG3’, nos2-forward 5’ GTTCTCAGCCCAACAATACAAGA3’, nos2-reverse 5’ GTGGACGGGTCGATGTCAC3’, chil3l3-forward 5’CAGGTCTGGCAATTCTTCTGAA3’, chil3l3-reverse 5’GTCTTGCTCATGTGTGTAAGTGA3’, arg1-forward 5’CTCCAAGCCAAAGTCCTTAGAG3’, arg1-reverse 5’AGGAGCTGTCATTAGGGACATC3’. The date generated was normalized to the lowest value.

### Statistical analysis

Statistical analysis was done using either Mann-Whitney U test to compare two groups or a one-way ANOVA to compare multiple groups. Values less than or equal to 0.05 were considered statistically significant. For RNA-seq analysis the false discovery rate (FDR) of less than 0.05 was used to determine differentially expressed genes and GO processes.

## Supplemental Figure Captions

**Supplemental Figure 1. Histology measurements of mouse tails**. C57BL/6J mice were infected with 2×10^6^ CFU of *Mm*. Beginning at day 7 post infection, mice were treated with imatinib (100mg/kg/day) or water for 7 days. Cross sections of the tail were cut and stained with H&E (A-N) or acid-fast bacillus (AFB) stain. **A**. Representative H&E images showing various features of the tail lesions scored in **B-N**. The upper two panels are at a magnification of 100x, and the lower two panels are at a magnification of 200x. **B-G**. Pathology scores for the epidermis and dermis for ulceration **(B)**, necrosis **(C)**, calcification **(D)**, edema **(E)**, acute inflammation **(F)**, and chronic inflammation**(G). H-L**. Pathology scores for the muscle for necrosis **(H)**, edema **(I)**, thrombi **(J)**, acute inflammation **(K)**, and chronic inflammation **(L). M**,**N**. Pathology scores for the bone for necrosis **(M)**, and inflammation **(N). O**. Representative images of AFB stain in the mouse tail lesions with micro-organisms identified in circles at a magnification of 1000x. Each data point represents scoring from one individual mouse in 5 separate experiments (n=21/group). Statistical test used was two-tailed Mann-Whitney U test, with p values did not reach significance (ns, not significant). The mean +/- SD for each group was graphed to show variance of data.

**Supplemental Figure 2. GO analysis of imatinib induced and infection related genes. A**,**B**. Genes differentially expressed with imatinib treatment in the absence of infection were identified by comparing differentially expressed genes from the uninfected water group, to imatinib-treated mice (514 genes, FDR < 0.05). Selection of GO terms identified by GO analysis of 373 genes upregulated with imatinib treatment (**A)**. Selection of GO terms identified by GO analysis of 141 genes downregulated with imatinib treatment (**B). C**,**D**. Selection of GO terms identified by GO analysis of the 903 genes upregulated at the 1 week infection timepoint **(C)** and the 676 genes upregulated at the 3 week infection timepoint **(D)** identified in Figure 3b. **E**. Comparison of **“**Infection Genes” (green circle; genes identified in Figure 3B), imatinib genes at 1 week infection (red circle; genes identified in Figure 3B), and imatinib genes at 3 weeks of infection (yellow circle; genes identified in Figure 3C). **F-H**. Comparison of 3 week infection genes identified in Figure 3B, to 3 week infection with imatinib genes identified in Figure 3C **(F)**. Selection of GO terms identified by GO analysis of the 814 upregulated genes **(G)**, and the 369 downregulated genes **(H)** specific to imatinib treatment at the 3 week infection time point.

**Supplemental Figure 3. Imatinib effects on BMDM during Mm infection. A**. RNA was collected from BMDMs derived from C57BL/6J mice infected with Mm at an MOI of 10 and left untreated or treated with 1μM imatinib for 8hrs, 24 hrs, or 44hrs. qPCR was used to measure levels of *nos2* in the cells at each time point (n= 2 wells/ group; data are representative of 3 separate experiments). **B-D**. BMDMs derived from C57BL/6J mice infected with Mm at an MOI of 10 and left untreated or treated with 1μM imatinib for 24 hrs. RNA was isolated from the cells and qPCR was used to measure levels of *nos2* **(B)**, *chil3l3* **(C)**, and *arg1* **(D;** n= 6 wells/ group). Statistical test used was one-way ANOVA, with p values indicated. The mean +/- SD, for each group was graphed.

