## Supplementary figures and images for "The host-directed therapeutic imatinib mesylate accelerates immune responses to *Mycobacterium marinum* infection and limits pathology associated with granulomas"

### Supplementary Figure 1

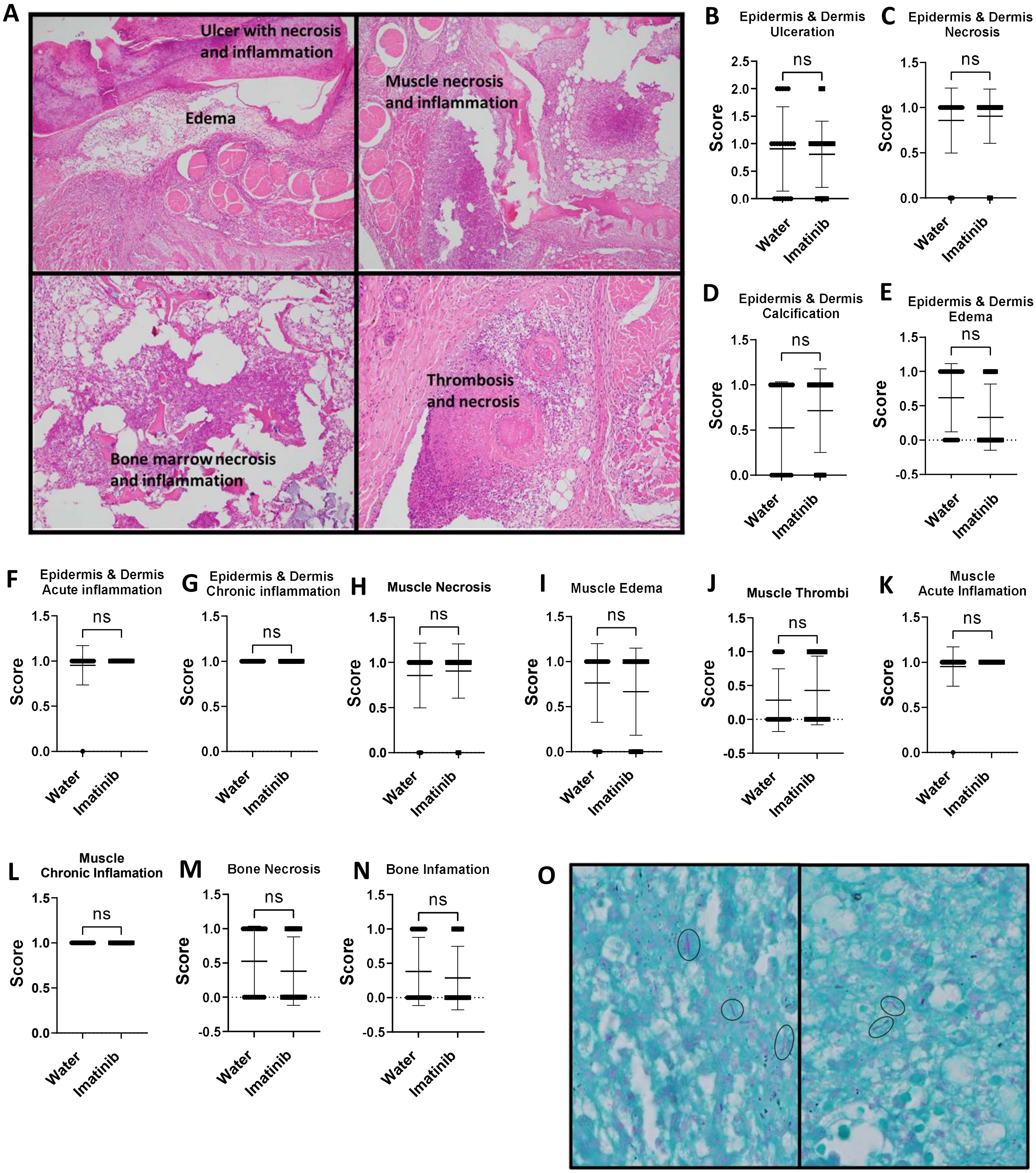


**Supplemental Figure 1. Histology measurements of mouse tails.**

### Supplementary Figure 2

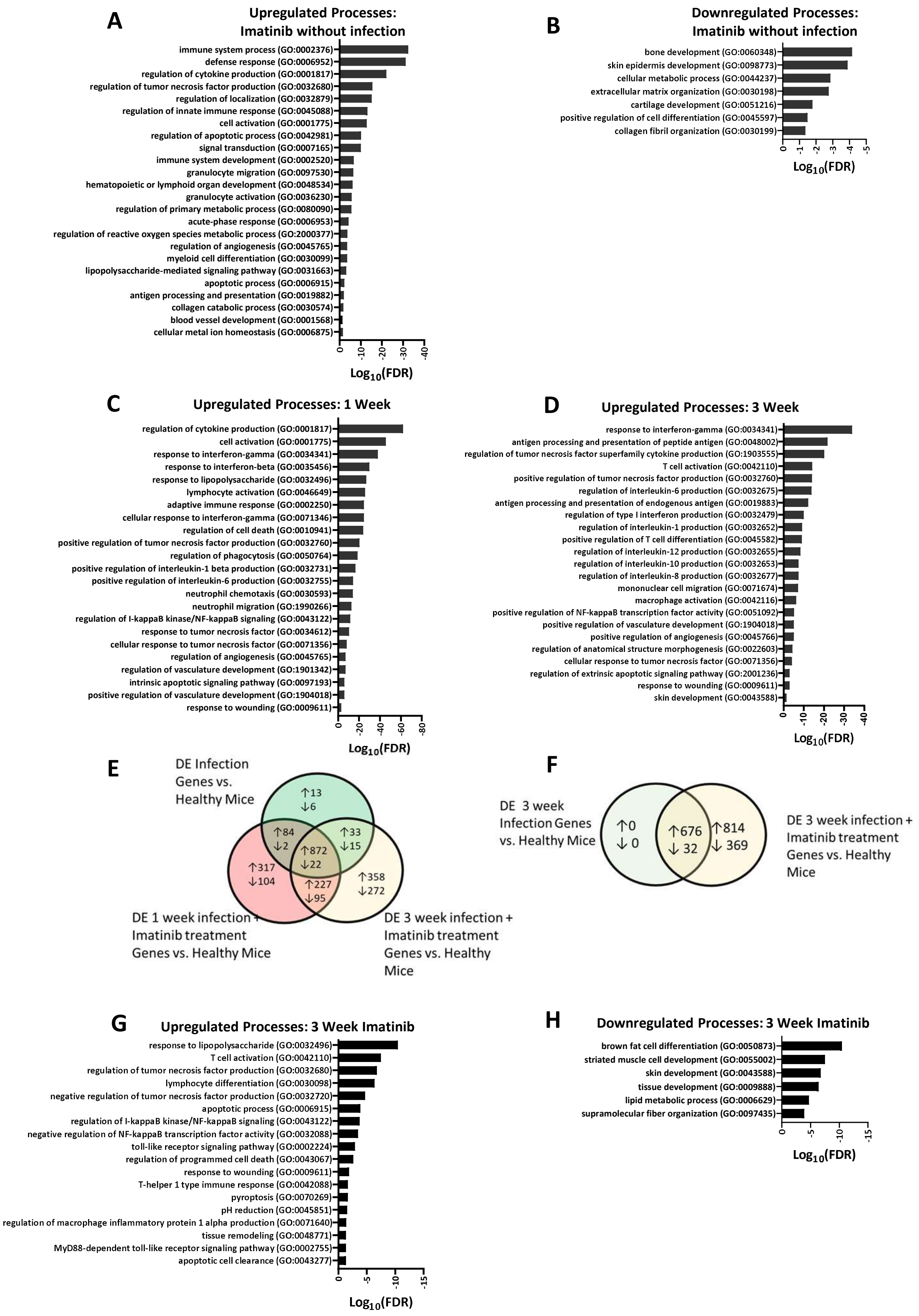


**Supplemental Figure 2. GO analysis of imatinib induced and infection related genes.**

### Supplementary Figure 3

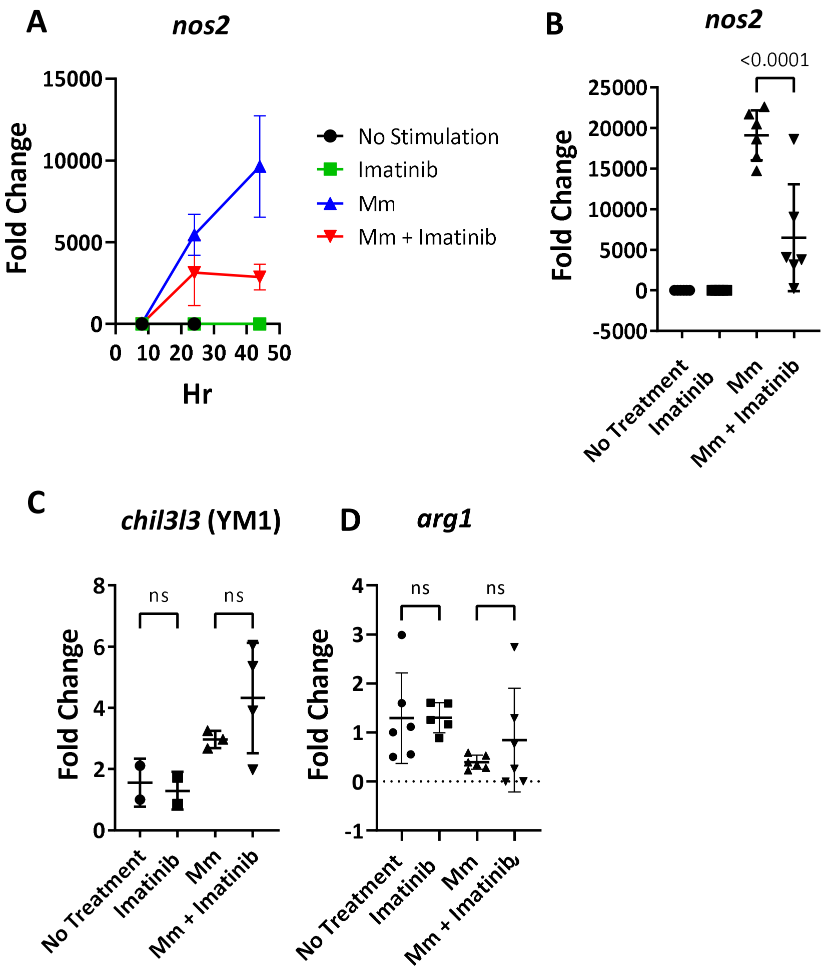


**Supplemental Figure 3. Imatinib effects on BMDM during Mm infection.**
